# TRAIL receptor agonist TLY012 in combination with PD-1 inhibition promotes tumor regression in an immune-competent mouse model of pancreatic ductal adenocarcinoma

**DOI:** 10.1101/2024.08.29.610345

**Authors:** Anna D. Louie, Kelsey E. Huntington, Young Lee, Jared Mompoint, Laura Jinxuan Wu, Seulki Lee, Thomas J. Miner, Wafik S. El-Deiry

## Abstract

Pancreatic ductal adenocarcinoma (PDAC) has an immunosuppressed, apoptosis-resistant phenotype. TLY012 is a pegylated recombinant Tumor necrosis factor-Related Apoptosis-Inducing Ligand (TRAIL), an orphan drug for chronic pancreatitis and systemic sclerosis. Innate immune TRAIL signaling suppresses cancer. We hypothesized that combination of immune checkpoint-blocking anti-PD-1 antibody and TLY012 would have synergistic anti-tumor efficacy in immune-competent PDAC-bearing mice. PDAC tumor-bearing C57Bl/6 mice treated 10 mg/kg anti-mouse PD-1 antibody twice weekly and 10 mg/kg TLY012 three times weekly had reduced tumor growth and tumor volume at 70 days compared to either drug alone (all p<0.005). B-cell activating factor (BAFF), which promotes PDAC tumors, decreased to 44% of control mice with dual treatment at 7 days and remained decreased at 3 months. Long-term dual treatment showed the highest levels of proinflammatory cytokines interferon gamma (average 5.6 times control level, p=0.046), CCL5 (average 14.1 times control level, p=0.048), and interleukin-3 (IL-3, average 71.1 times control level, p=0.0053). Flow cytometry showed trends toward decreased circulating regulatory T cells, increased NK cells, and a higher proportion of CD8+ T cells within tumors in dual treatment group. In summary, combination of anti-PD-1 and TLY012 prevented growth of PDAC in an immunocompetent mouse model while increasing tumor-infiltrating CD8+ T cells, decreasing circulating T-regulatory cells and altering cytokine expression of CCL5, interferon gamma and IL-3 to promote proinflammatory, antitumor effects. Combining TLY012 and anti-mouse PD-1 creates changes in immune cell and cytokine levels to induce a more proinflammatory immune environment that contributes to decreased PDAC tumor growth.

## Introduction

Pancreatic ductal adenocarcinoma (PDAC) remains a deadly disease with little progress in long-term survival over the last several decades [1-5]. Much work has been done to characterize molecular alterations and drivers in pancreatic cancer including among different ethnic groups, early onset pancreatic cancer, and treatment effects on genomic landscape [6-13]. Some of the key driver alterations include mutations in KRAS, p53, p16, SMAD4, KDM6A, ARID1A, and BRCA2, signatures of DNA damage and repair deficiency, mismatch repair deficiency as well as amplified genes such as ERBB2, MET, PIK3CA among others [6, 7, 11, 12]. While much progress has been made with immunotherapy across tumor types, and much understanding of the biology of pancreatic cancer, there has been little impactful progress on advanced PDAC in part due to an immunosuppressed, apoptosis-resistant phenotype [14-23].

The TNF-Related Apoptosis-Inducing Ligand (TRAIL) is part of the host immune system that suppresses transformed cells, virally infected cells as well as cancer and its metastases [24-35]. Nearly 3 decades ago, the discovery of pro-apoptotic TRAIL Death Receptor DR5 as a direct transcriptional target of the p53 tumor suppressor gene directly linked the innate immune system to the host response to suppress tumorigenesis [36-42]. Deletion of TRAIL receptor DR5 in mice led to apoptotic resistance in vivo as well as tumorigenesis [43, 44]. It was later recognized that the TRAIL gene is also a p53 target gene and subsequent work identified TRAIL-inducing compound #10 (TIC10; also known as ONC201) [45, 46]. We previously reported that the combination of TRAIL or TRAIL receptor DR5 agonist antibodies and ONC201 has particularly potent anti-tumor effects *in vivo* across cancer types including pancreatic cancer [47-50]. We have also previously reported a TRAIL-inducing micro-RNA that we have not, as of the date of publication of this manuscript, translated to *in vivo* studies or to treatment of patients [51].

The TRAIL pathway influences both inflammation and tumorigenesis as well as fibrosis, all relevant to PDAC and its potential novel therapeutics [44, 52-55]. TLY012 is a pegylated recombinant Tumor necrosis factor-Related Apoptosis-Inducing Ligand (TRAIL) [55, 56].

In the present studies we explore the hypothesis that TLY012’s modulation of the tumor microenvironment has potential for synergistic effects when combined with immune checkpoint blockade. We demonstrate potent anti-tumor effects of TLY012 combined with anti-mouse PD1 and demonstrate immune alterations including in B-and T-cell immunity as well as tumor promoting cytokines. Our results prompt further preclinical and clinical studies evaluating the novel treatment combination of TLY012 and anti-PD1.

## Materials and Methods

### Cell Culture

Cell lines used for this study were acquired from the American Type Culture Collection (ATCC), unless otherwise indicated. All pancreatic cancer cells were grown in Dulbecco’s Modified Eagle Medium (DMEM) with 10% Fetal Bovine Serum (FBS). Human Foreskin Fibroblasts (HFF) were grown in DMEM with 15% FBS. rhTRAIL was generated in-house using a protocol previously developed by our lab and detailed in *Kim et al*. TLY012 was provided by Theraly Fibrosis, Inc. TLY012 was diluted in sterile phosphate buffered saline (PBS) for *in vitro* experiments.

### Cell viability Assay

To assess cell viability, cells were plated overnight in 96-well plates at a density of 1.0 × 10^4^ cells/well. All cells were plated in triplicates. After 72 hours, the media in the wells was replaced with either fresh media (controls) or with media containing various doses of rhTRAIL or TLY012. After 4 hours of incubation, cells were treated with Cell-Titer Glo (Promega) and imaged to assess cell viability. Synergy and combination indices were determined using Compusyn, which uses the Chou-Talalay method for determining synergy.

### Flow Cytometric Analysis

Sub-G1 analysis was conducted following a well-established lab protocol. Cells were plated overnight before treating with drugs. After the experiment was completed, all floating and adherent cells were harvested. Once cells were harvested, they were gently centrifuged and washed with PBS to remove the phenol red found in the culture medium. Once thoroughly washed, cells were resuspended and fixed in 70% cold ethanol and stored at 4°C overnight. Once cells were fixed, they were stained with propidium-iodide to bind DNA. Cells were then run on an Epics Elite (Beckman Coulter) flow cytometer. FlowJo was used to analyze the percentage of apoptotic cells.

### Western Blotting

Cells were plated overnight before any drug treatment. Approximately 5 × 10^5^ cells/well were plated in 6-well plates. Upon completion of the experiment, 1 mL of media containing floating cells was harvested. Adherent cells were crushed and collected into the remaining 1 mL of media, and this total solution was pelleted for 5 minutes at 400 Rcf. The pellets were then washed by resuspending with PBS once. Cells were then lysed using RIPA buffer (Sigma-Aldrich) with 1X protease inhibitor and 1x phosphatase inhibitor (Roche). Upon lysis, cells were centrifuged at 13,000 RPM at 4°C for approximately 20 minutes. The supernatants were collected. Proteins were estimated using a BCA Assay (Thermo Scientific). Gels used for analysis were pre-cast NuPAGE 4-12% Bis-Tris (Thermo Scientific). Bands were quantified using an established ImageJ protocol from the ImageJ user guide.

### *In Vivo* Tumor Xenograft Studies

All *in vivo* studies conducted for this manuscript were approved by the Brown University IACUC. For *in vivo* tumor xenograft studies, we used female C57Bl/6 mice. Mice were aged 5-8 weeks at the time of tumor inoculation. Cells were mixed in a 50:50 Matrigel (Corning):PBS solution and mixed at various dilutions. Total inoculation volume was 200 µL, irrespective of tumor model or number of cells inoculated. The vehicle is a solution of 20% Cremophor EL (Sigma-Aldrich), 70% PBS, and 10% DMSO. rhTRAIL was administered through intravenous tail vein injections. TLY012 was administered via intraperitoneal injections.

### Tumor Volume Calculations

Measurements were taken using Vernier calipers. Thus, the equation for calculating tumor volume is: Volume = (Width^2^ * Length)/2. Treatment was initiated once the tumors reached an optimal volume between 100-150mm^3^.

### Immunohistochemical Staining

Tumors were fixed in formalin immediately after harvesting in cassettes. After fixation, cassettes were paraffin embedded. Slides were cut 5□m thick. Immunohistochemistry was initiated by deparaffinizing slides using xylene. Slides were dehydrated through sequential dilutions of ethanol. The antigen retrieval step was conducted by heating slides for 10 minutes in pH 6.0 citrate acid buffer. Ki67 (MIB-1) antibody was obtained from Cell Signaling Technologies, used at 1:200 dilution. CC3 Antibodies obtained from BD Biosciences, used at 1:100 dilution. Slides were incubated in primary antibodies overnight; respective secondary antibodies were added the following day. Slides were developed using DAB Staining Kit (Vector Labs) and mounted using a xylene-based mount, Cytoseal XYL.

### Immunohistochemical quantification

Slides were quantified using QuPath, an open-source, automated program for immunohistochemistry quantification.^26^ Ki67 staining was analyzed in a binary fashion with only positive and negative nuclei. CC3 staining was analyzed by taking 5 representative high power (20x) fields per tumor and using QuPath to quantify cells that stained positive.

### Statistical Analysis

All statistical analyses were conducted using Microsoft Excel and figures were designed using Prism GraphPad. P-values were determined using the Student’s T-test and significance was determined as P < 0.05.

## Results

### Potent anti-tumor effects in PDAC-implanted syngeneic C57Bl/6 mice treated with anti-mouse PD-1 and TLY012

TLY012 taps into the host innate immune system and targets fibrosis within the tumor microenvironment. These properties including removal of an immune suppressive microenvironment are well suited for synergy with other agents such as immune checkpoint therapy that may have improved penetration and efficacy in the tumor microenvironment.

We set up a mouse model treatment scheme (**Figure 1A**) to test the hypothesis that combination of immune checkpoint-blocking anti-PD-1 antibody and TLY012 would have synergistic anti-tumor efficacy in immune-competent PDAC-bearing mice. The results of the long-term *in vivo* study demonstrates potent anti-tumor effects in PDAC-implanted syngeneic C57Bl/6 mice treated with anti-mouse PD-1 and TLY012 (**Figure 1B**). Available tumors at the end of the experiment were photographed (**Figure 1C**). The mice tolerated the treatments well with no reduction in mouse weights (**Figure 1D**).

**Figure 1.**
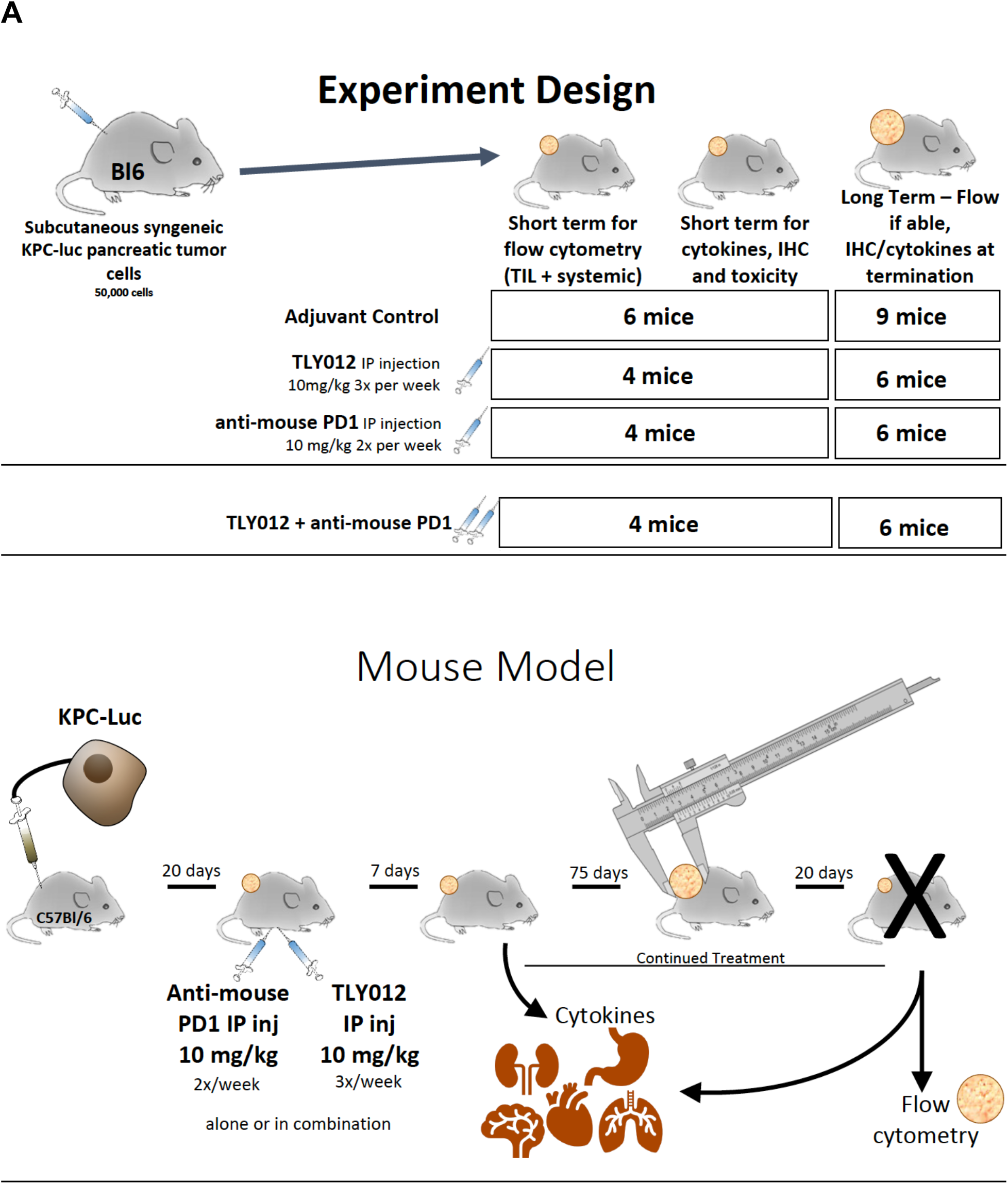
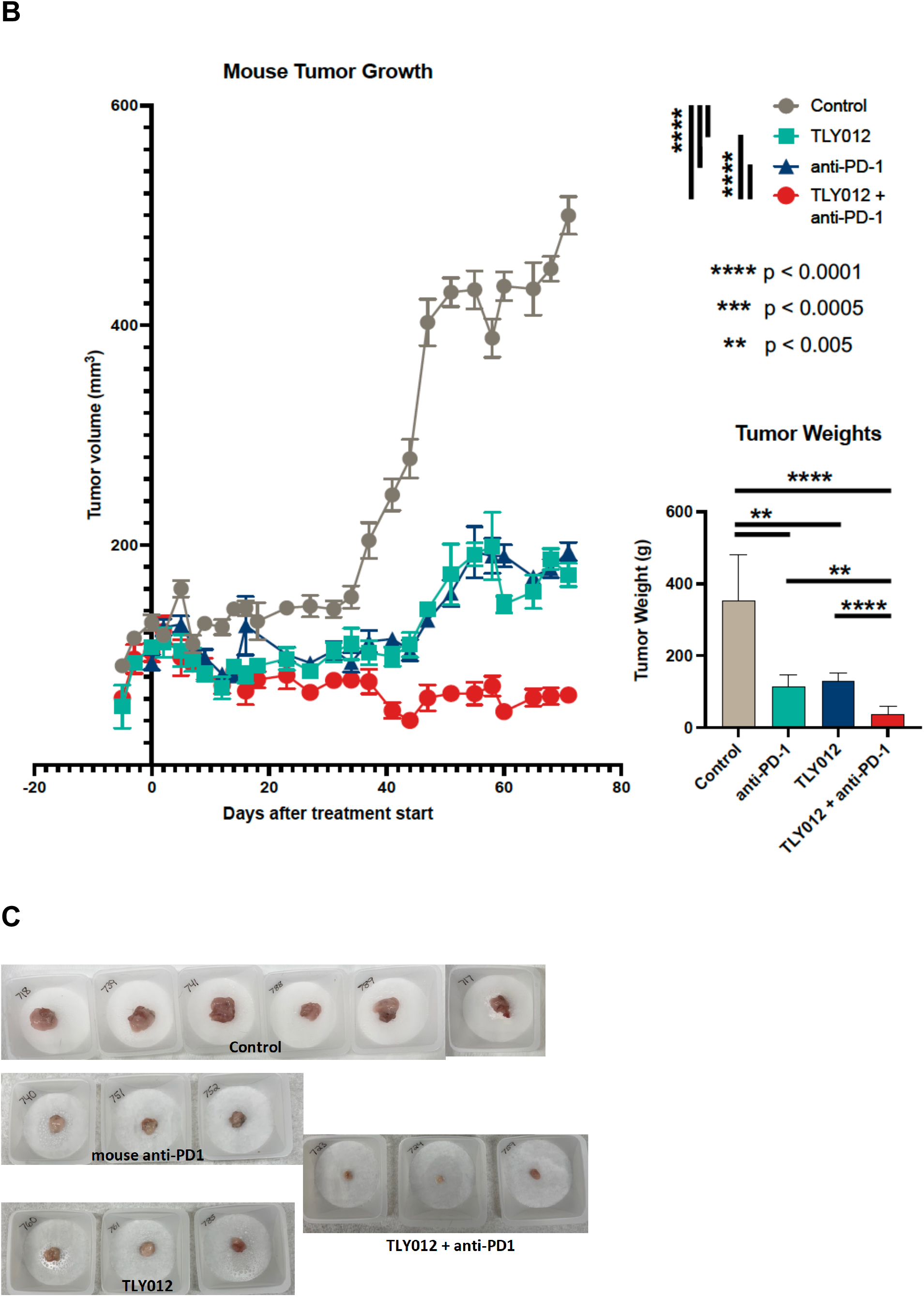
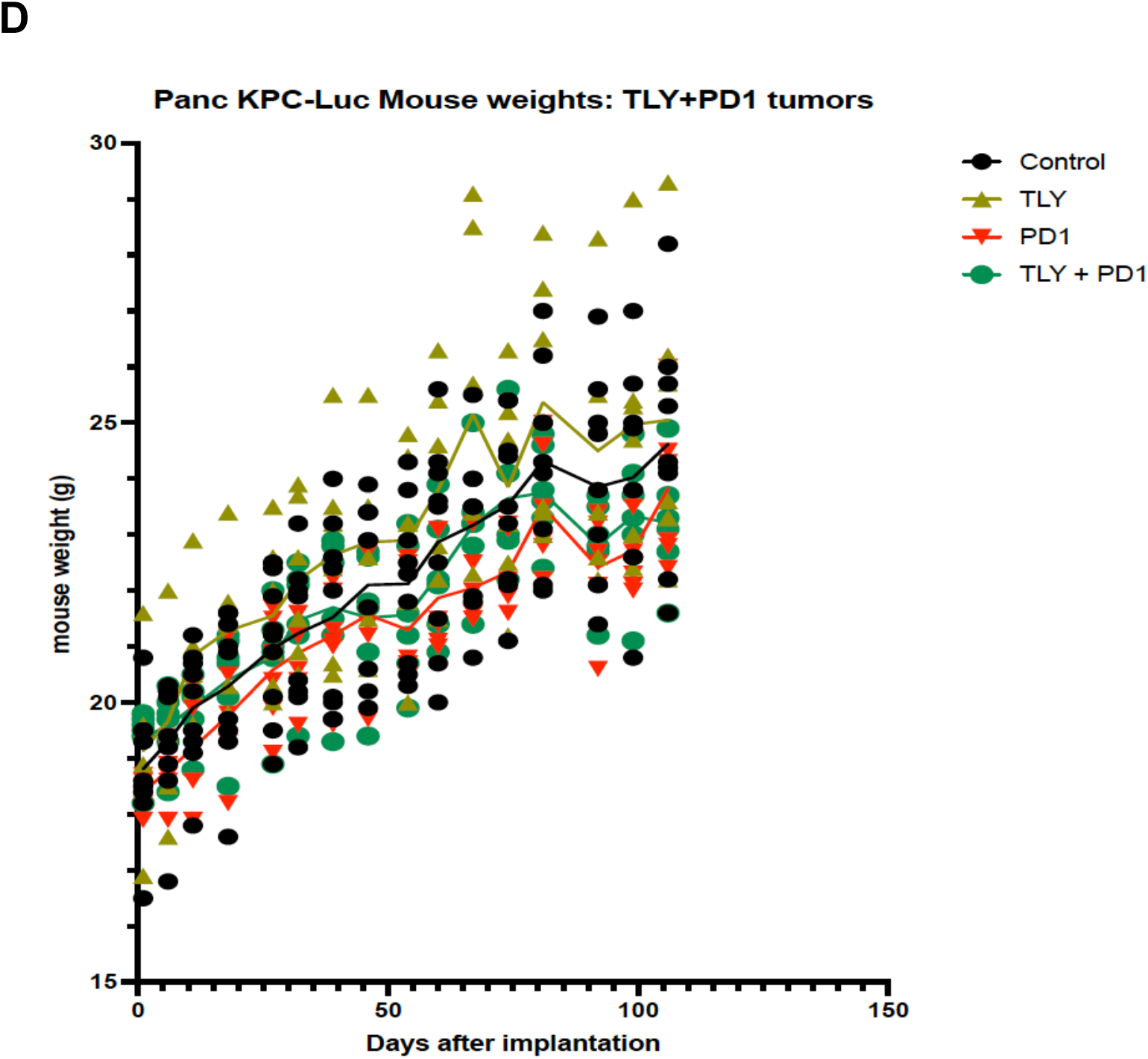
Syngeneic mice with PDAC treated with both anti-mouse PD-1 and TLY012 delays tumor growth and reduces tumor volume. **(A)** *In vivo* PDAC mouse model and treatment schema for anti-tumor efficacy, biomarker, and immune analysis. The number of mice used for short-term as well as long-term treatments is as indicated for each cohort. **(B)** Syngeneic mice with PDAC treated with both anti-mouse PD-1 and TLY012 resulted in slower tumor growth and reduced tumor volume at 70 days compared to either drug alone (all p<0.005). Tumor weights are shown in the graph in the lower right for the respective treatment groups. **(C)** Images of available tumors at the end of experiment are shown. **(D)** Weights of mice in different treatment groups over the course of the experiment.

### Decreased circulating regulatory T-cells, increased NK-cells, and a higher proportion of CD8+ T cells within tumors after dual treatment with anti-mouse PD-1 and TLY012

We performed flow cytometric analyses of spleen and tumor T-cells and NK-cells to examine the effects of individual anti-mouse PD-1, TLY012 as well as the dual anti-mouse PD-1 and TLY012 treatment group in treated syngeneic C57Bl/6 KPC-Luc PDAC tumor-bearing mice (**Figure 2**).

**Figure 2.**
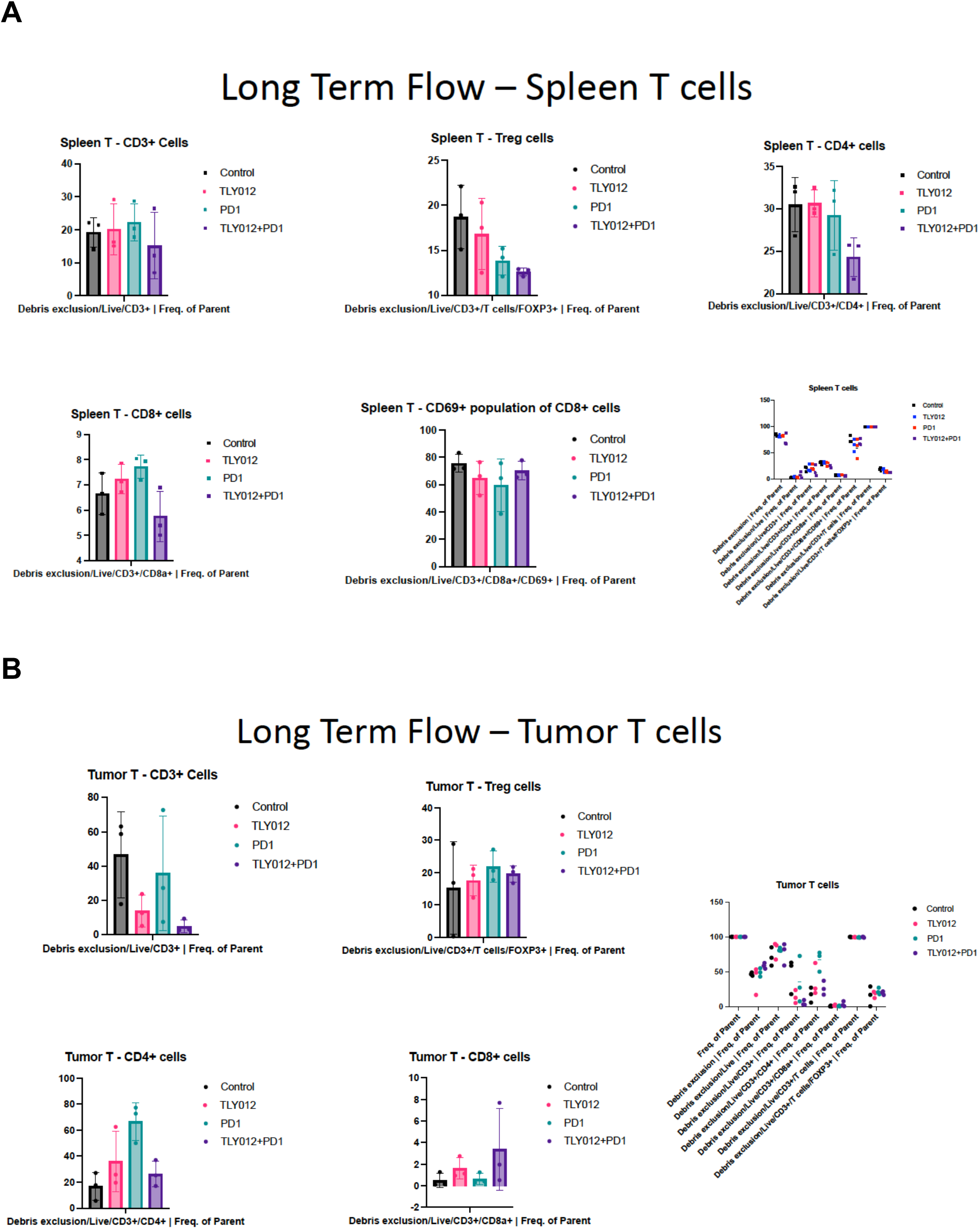
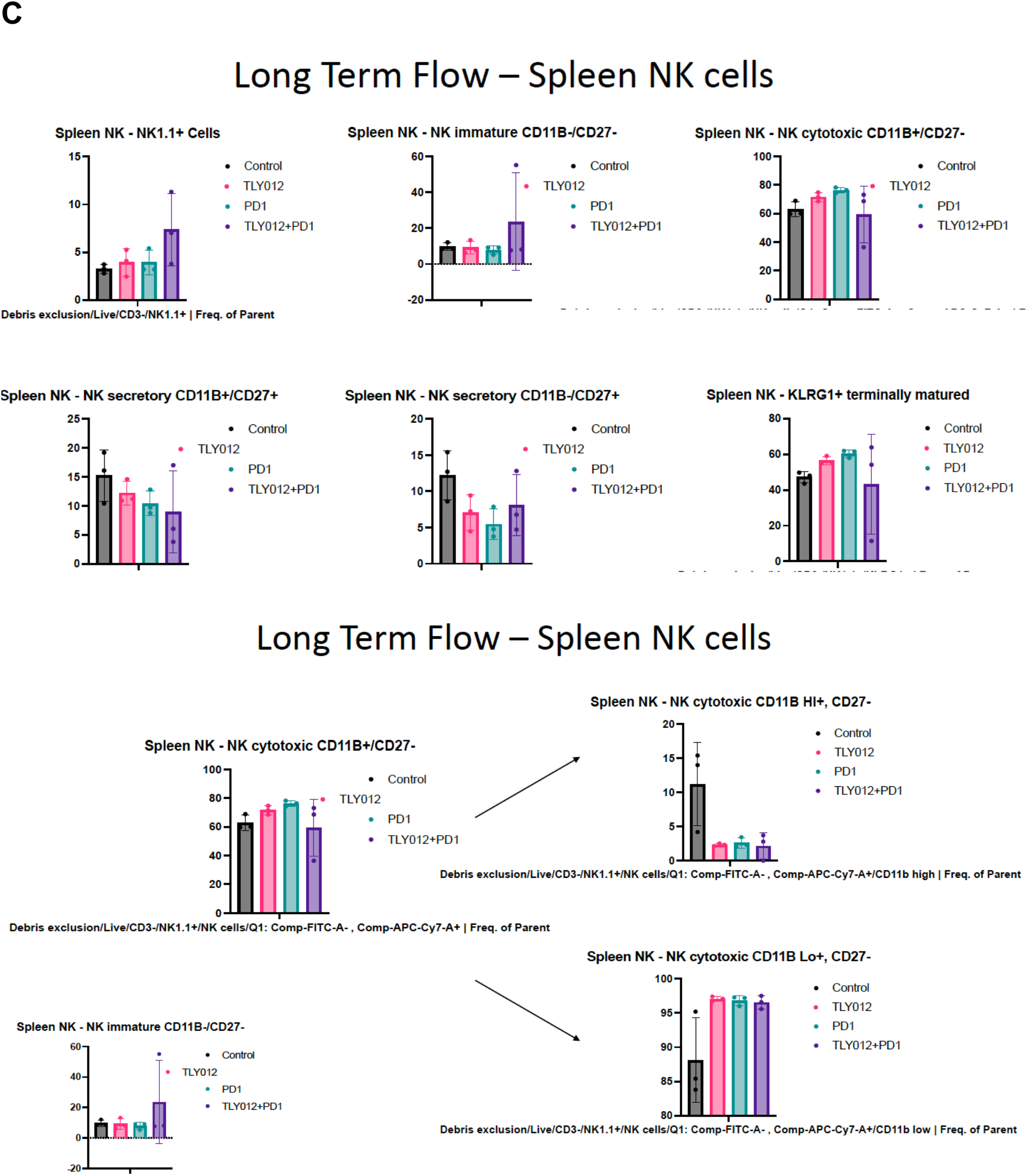
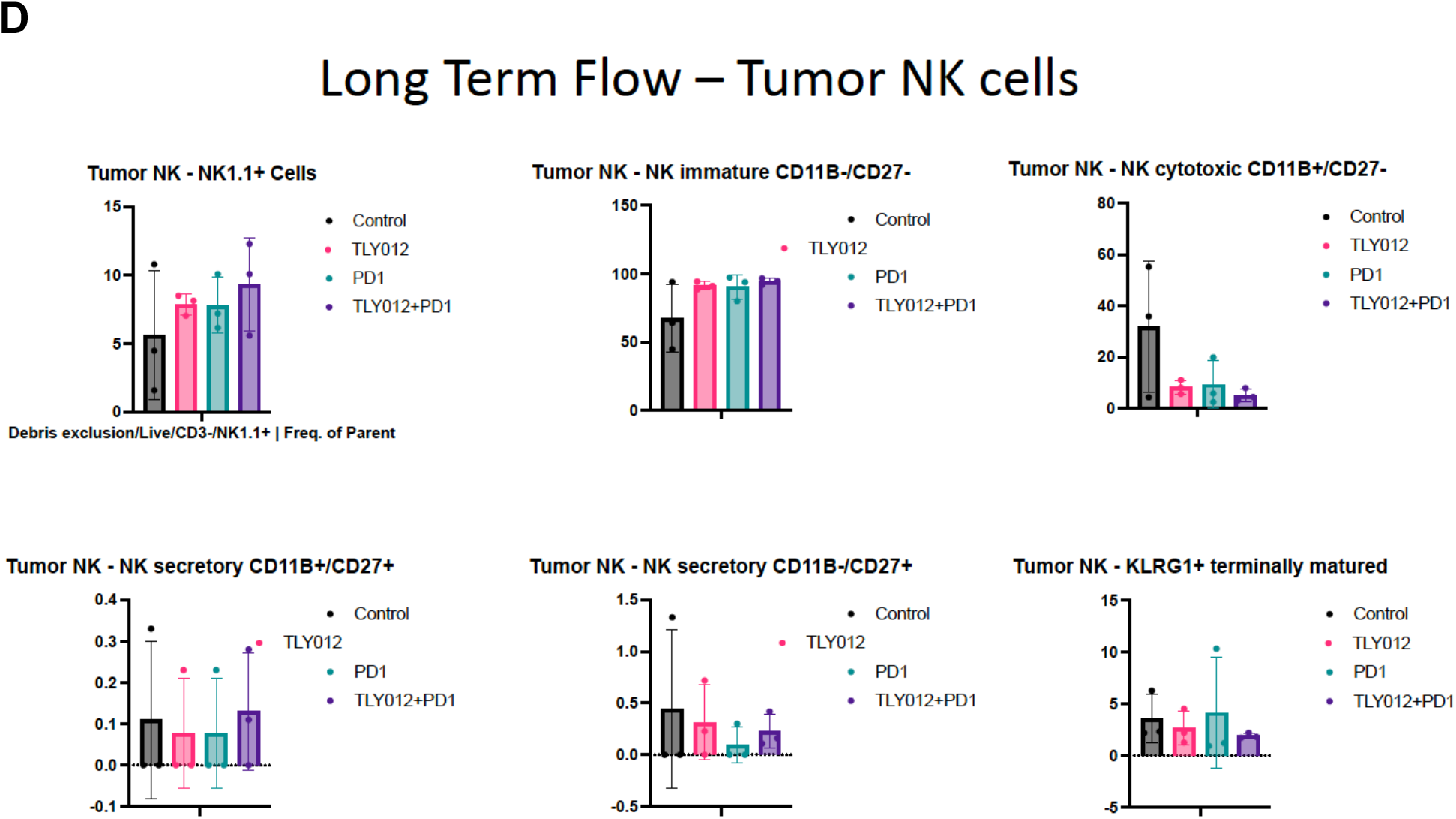
Decreased circulating regulatory T-cells, increased NK-cells, and a higher proportion of CD8+ T cells within tumors after dual treatment with anti-mouse PD-1 and TLY012. **(A)** Flow cytometric immune analysis of spleen T cells following treatment with TLY012, anti-PD1 or the combination of TLY012 and anti-PD1 in a syngeneic pancreatic cancer mouse model (KPC-Luc). **(B)** Flow cytometric immune analysis of tumor T following treatment with TLY012, anti-PD1 or the combination of TLY012 and anti-PD1 in a syngeneic pancreatic cancer mouse model (KPC-Luc). **(C)** Flow cytometric immune analysis of spleen NK cells following treatment with TLY012, anti-PD1 or the combination of TLY012 and anti-PD1 in a syngeneic pancreatic cancer mouse model (KPC-Luc). **(D)** Flow cytometric immune analysis of tumor NK cells following treatment with TLY012, anti-PD1 or the combination of TLY012 and anti-PD1 in a syngeneic pancreatic cancer mouse model (KPC-Luc).

We observed a reduction in splenic T-regulatory as cytotoxic CD8+ T-cells (**Figure 2A**) along with a relative enrichment in analyzed tumors of cytotoxic CD8+ T-cells in combination anti-mouse PD-1 and TLY012-treated tumors (**Figure 2B**) in the syngeneic KPC PDAC model. We observed enrichment of immature NK cells in the spleen of combo treated mice (**Figure 2C**) and similarly the tumors of dual anti-mouse PD-1 and TLY012 treated PDAC tumor-bearing C57Bl/6 mice showed enrichment with immature NK cells (**Figure 2D**). Thus, the combination of TLY012 plus anti-PD1 enriches treated tumors with cytotoxic CD8+ T-cells as well as immature NK cells. These findings are consistent with the observed synergistic anti-tumor efficacy of the combination therapy *in vivo*.

### B-cell activating factor (BAFF), which correlates with tumor progression in PDAC, was reduced in treatment groups at 7 days and 3 months

B-cell activating factor (BAFF) correlates with tumor progression in PDAC [57]. BAFF was reduced in treatment groups in anti-PD1 treated and dual TLY012 plus anti-PD1 treated mice at 7 days and was reduced in single and dual treated groups after three months (**Figure 3**). Reduced BAFF levels are consistent with the observed anti-PDAC effects of treatment with TLY012 plus anti-PD1.

**Figure 3.**
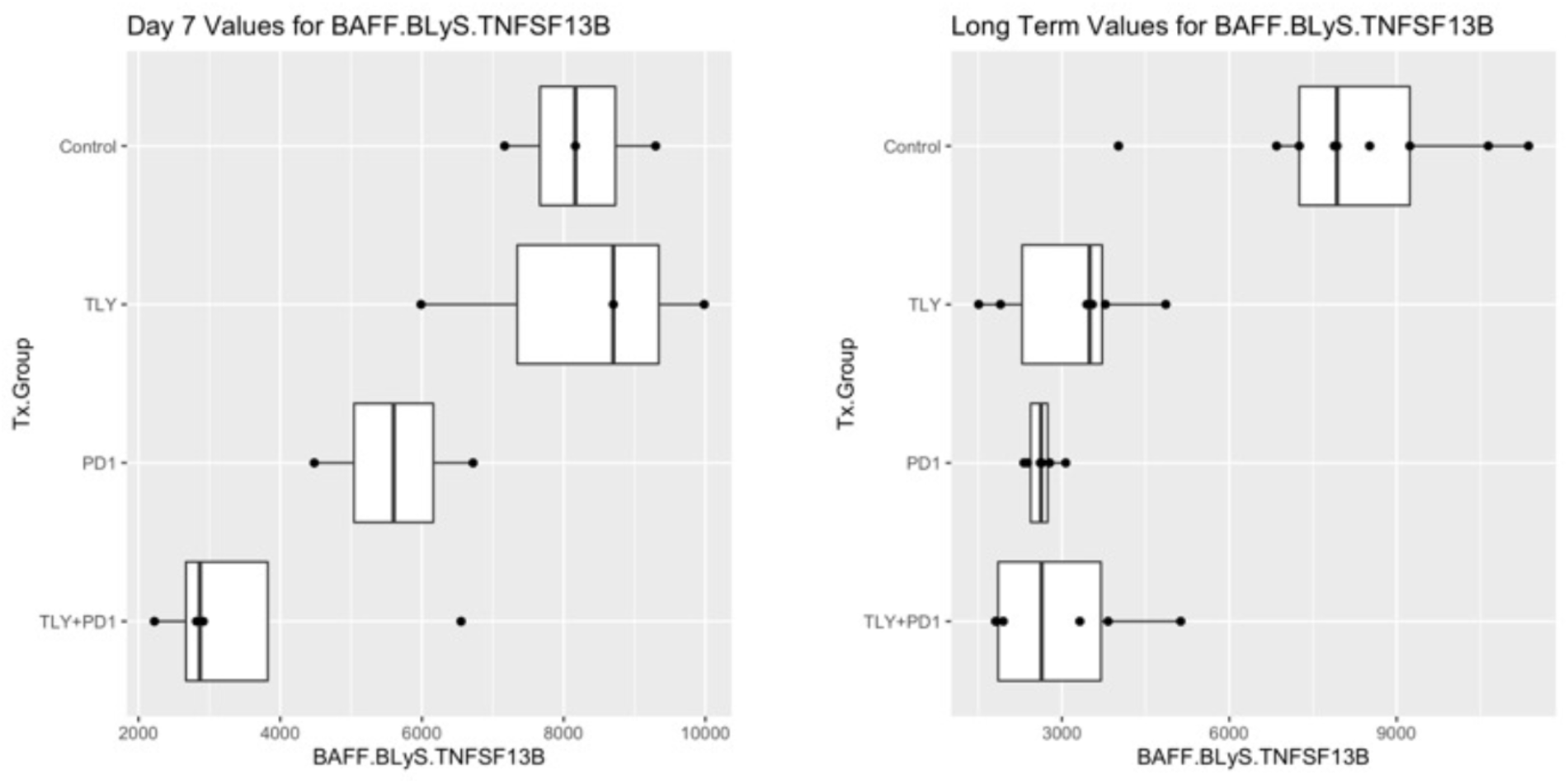
Reduced tumor promoting B-cell activating factor (BAFF) levels in in anti-PD1 treated and dual TLY012 plus anti-PD1 treated mice at 7 days and in single and dual treated groups at three months. Day 7 cytokine levels are in the left panel and the levels at 3 months are shown in the right panel for the different treatment groups as indicated. The numbers shown for serum BAFF levels on the X-axes are in pg/mL.

### Long-term serum cytokine analysis following treatment with TLY012, anti-PD1 or the combination of TLY012 and anti-PD1 in syngeneic KPC PDAC mouse model

We assessed serum cytokine levels at 3 months in the TLY012, anti-PD1 or combination treatment groups versus control mice (**Figure 4**).

**Figure 4.**
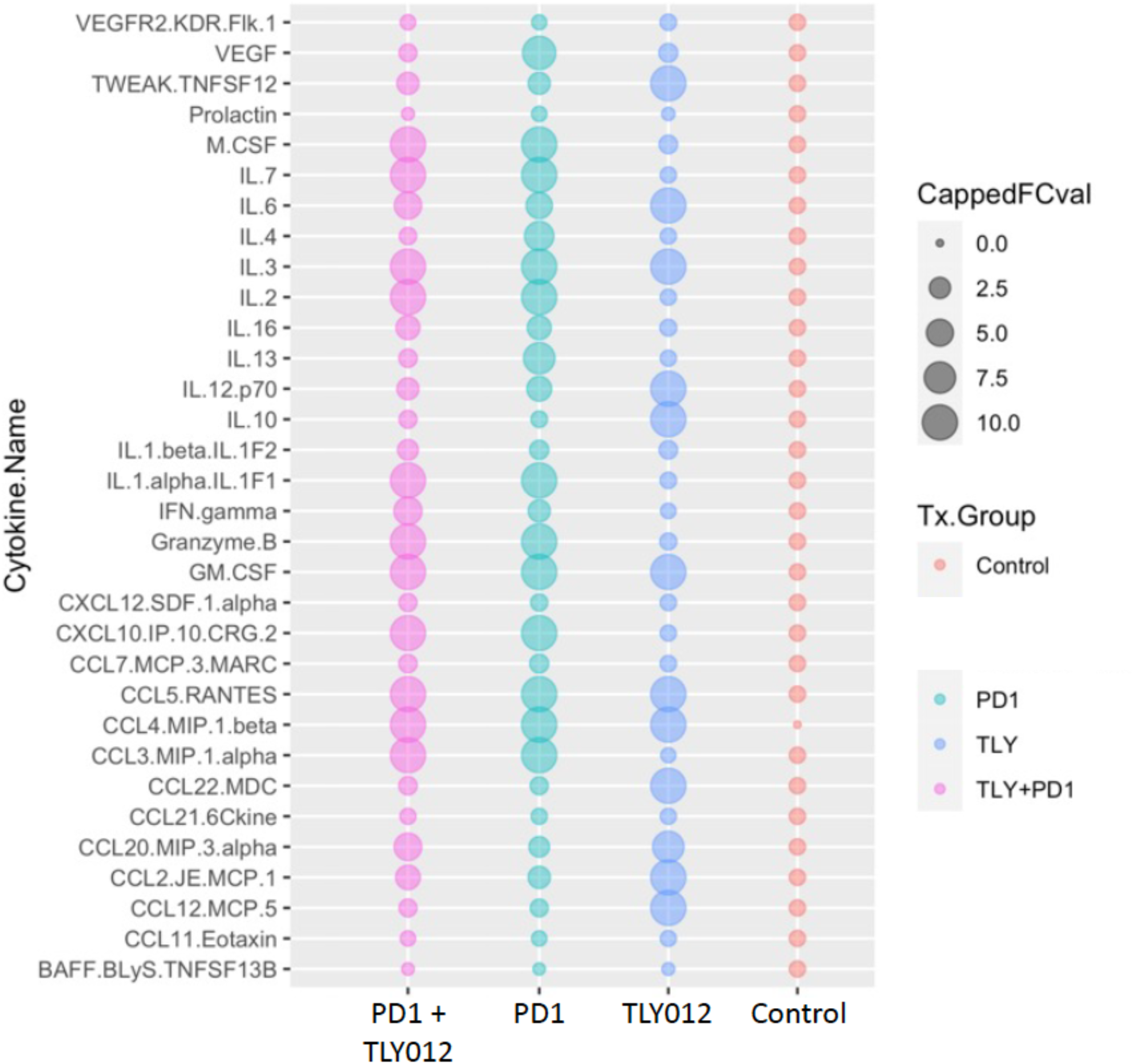
Long-term cytokine levels in KPC PDAC tumor-bearing C57Bl/6 mice for control, TLY012, anti-PD1, or dual therapy with TLY012 plus anti-PD1 as indicated.

A number of proinflammatory cytokines including interferon gamma and CCL5 were elevated while VEGFR2 trended towards reduction in the long-term dual treatment group (**Figure 4, 5**). These results are consistent with the observed anti-PDAC effects of treatment with TLY012 plus anti-PD1.

**Figure 5.**
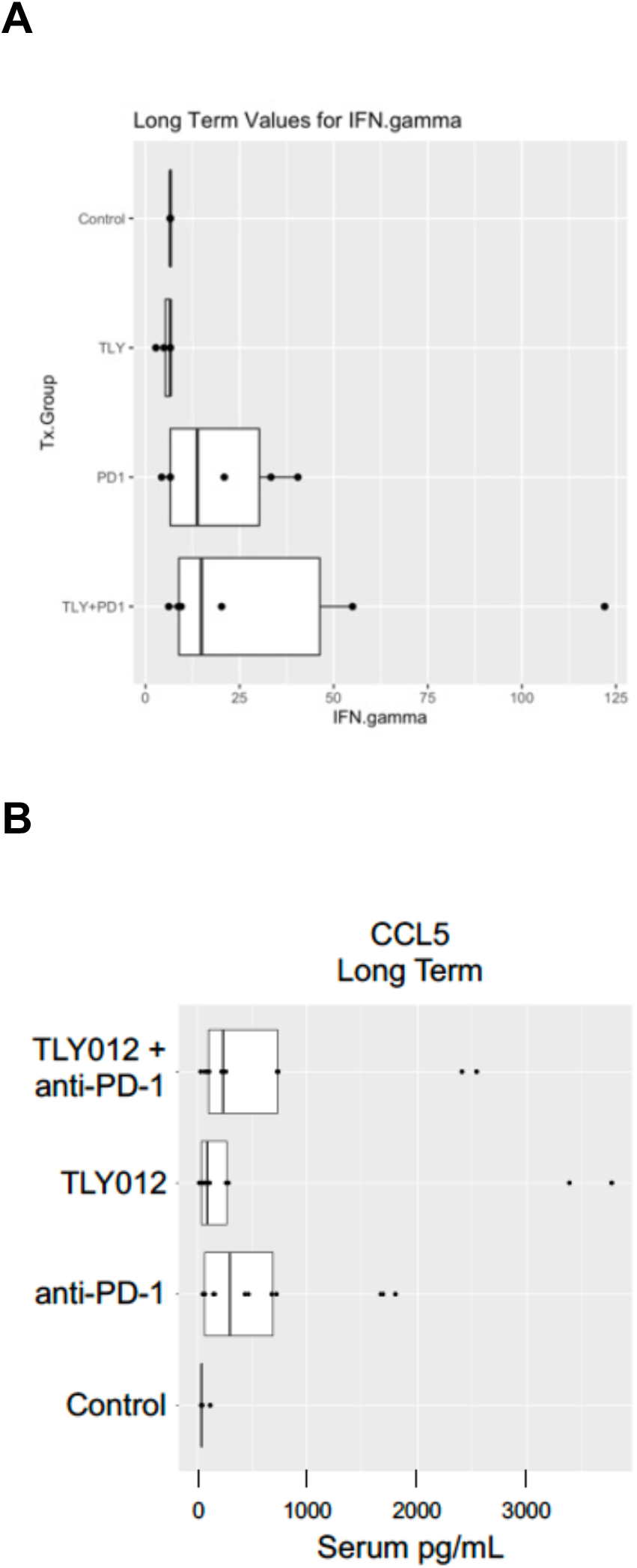
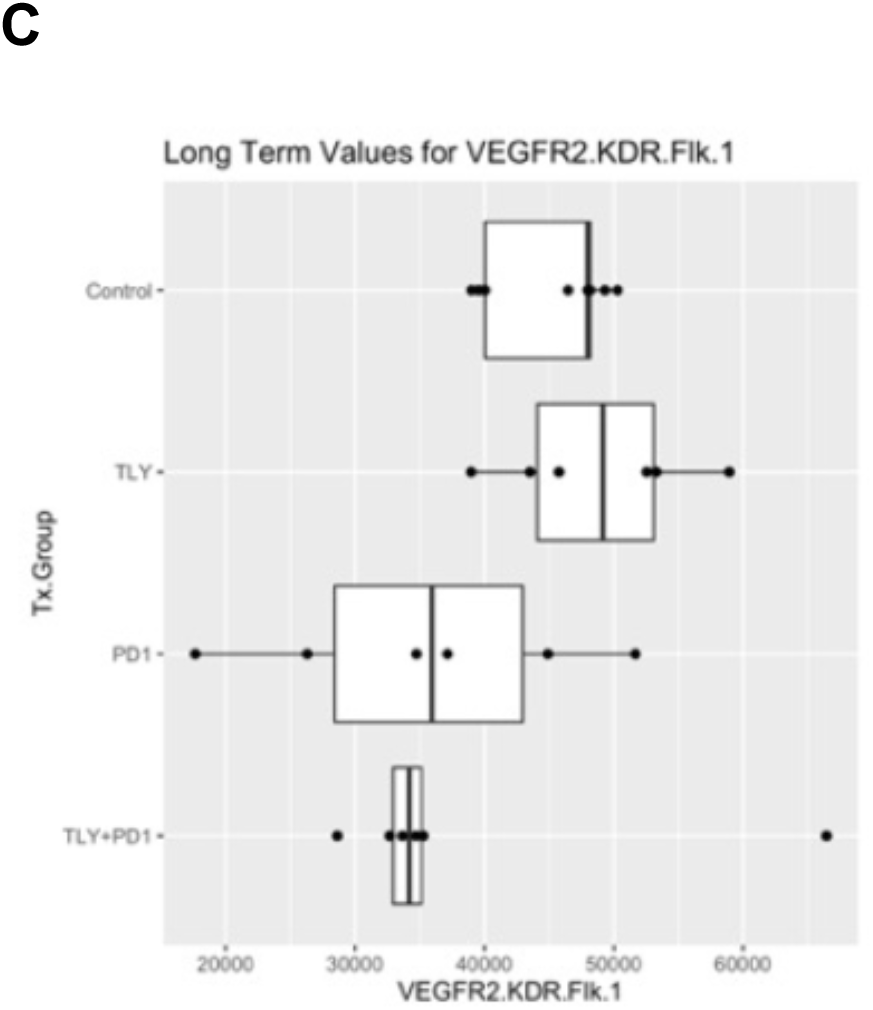
Alterations in selected cytokines at 3 months in KPC PDAC tumor-bearing C57Bl/6 mice for control, TLY012, anti-PD1, or dual therapy with TLY012 plus anti-PD1 as indicated. (A) IFN-gamma levels at 3 months in pg/mL for control, TLY012, anti-PD1, or dual therapy with TLY012 plus anti-PD1 as indicated. (B) CCL5 levels at 3 months in pg/mL for control, TLY012, anti-PD1, or dual therapy with TLY012 plus anti-PD1 as indicated. (C) VEGFR2 levels at 3 months in pg/mL for control, TLY012, anti-PD1, or dual therapy with TLY012 plus anti-PD1 as indicated.

## Discussion

We report potent anti-tumor effects following long-term treatment for 3 months with combination of innate immune anti-fibrotic TLY012 and immune checkpoint blocker anti-mouse PD-1 in a KPC PDAC syngeneic C57Bl/6 mouse model. The treatments were associated with changes in immune cell populations in spleens and tumors and serum cytokine levels to induce a more proinflammatory immune environment that correlates to decreased PDAC tumor growth.

A limitation of our study is that the tumors were implanted subcutaneously and not orthotopically which could influence the results due to differences in local microenvironment. Clearly this would need to be further addressed in future studies.

Another limitation is that while we performed flow cytometric analysis of tumor tissue and spleens, as well as serum cytokine profiling, we did not carry out immunohistochemical staining of tumors at early or late time points to provide further evidence of altered biomarkers and proposed mechanisms. We also did not test the requirement of NK-or T-cell alterations or cytokine alterations with regard to the observed anti-PDAC effects *in vivo*. We also did not evaluate the anti-fibrotic effects of TLY012 as monotherapy or in combination with anti-PD1 *in vivo*.

Despite the limitations, our results provide a novel preclinical combination therapy of innate immune TLY012 and immune checkpoint blocker anti-mouse PD-1 that has potent efficacy in a KPC PDAC syngeneic C57Bl/6 mouse model and without evidence of toxicity. The results support the further exploration and clinical testing of TLY012 alone and in combination with anti-PD1 therapy in pancreatic cancer as well as other malignancies.

## Acknowledgements

W.S.E-D. is an American Cancer Society Research Professor and is supported by the Mencoff Family University Professorship at Brown University. This work was supported by a grant from D&D Pharmatech to W.S.E-D. This work was presented in part at the 2022 meeting of the American Association for Cancer Research and the 2022 meeting of the Society of Surgical Oncology.

